# Single-cell dynamics of genome-nucleolus interactions captured by nucleolar laser microdissection (NoLMseq)

**DOI:** 10.1101/2024.06.18.599512

**Authors:** Kaivalya Walavalkar, Shivani Gupta, Jelena Kresoja-Rakic, Mathieu Raingeval, Chiara Mungo, Raffaella Santoro

## Abstract

Gene position in the nuclear space plays an important role in gene regulation. This is exemplified by repressive chromatin domains frequently contacting nuclear lamina or nucleoli. The nucleolus undergoes structural changes in response to various cellular states, potentially impacting genome organization. However, how the 3D-genome organization responds to nucleolar states has remained underinvestigated due to the lack of methods able to identify nucleolar associated domains (NADs) in single cells and under nucleolar stress. To address this, we developed NoLMseq, a method combining laser-capture microdissection and DNA sequencing to map NADs in single cells. NoLMseq identified many unexplored features of chromosome organization around single nucleoli such as NAD heterogeneity among ESCs, culminating in two major populations with distinct chromatin states. NADs prevalently contact nucleoli in a monoallelic manner and allelic nucleolar contact frequency mirrors gene expression and chromatin states. NoLMseq also revealed how chromosomes reorganise around nucleoli under nucleolar stress conditions, highlighting the importance of nucleolus integrity in genome organization and 3D-genome response to nucleolar stress. The results demonstrated that NoLMseq accurately measures chromosome contacts around single healthy and stressed nucleoli and it will be a critical tool to study NADs within biological populations and determine how the 3D-genome responds to nucleolar stress in healthy and disease states.

## Introduction

The three-dimensional (3D) organization of chromosomes in the cell’s nucleus plays an important role in the regulation of gene expression programs and cell fate. One important aspect of this regulation is the position of genes in the nuclear space. This is exemplified by the frequent location of repressive chromatin domains at the nuclear periphery (i.e., lamina associated domains, LADs) or around nucleoli (nucleolar associated domains, NADs) ^1–3^. However, while LADs have extensively been studied in many cell types and in single cells, NADs have remained underinvestigated mainly due to technical limitations in identifying DNA sequences associating with a compartment lacking a membrane such as the nucleolus.

The nucleolus exhibits a highly dynamic structure that changes in response to external stimuli affecting ribosome biogenesis, a critical nucleolar process that is tightly regulated according to cell state ^4^. These nucleolar structural alterations (i.e., size, number, or physico-chemical properties) might significantly impact the organization of the surrounding genome, thereby affecting gene expression and chromatin states. Ribosome biogenesis is initiated in the nucleolus by the RNA polymerase I (Pol I)-driven transcription of hundreds of ribosomal RNA (rRNA) genes (rDNA) that generate 45S/47S pre-rRNA. These transcripts are then modified, processed, and assembled with ribosomal proteins (RPs) in the nucleolus, forming pre-ribosomal particles ^5^. rRNA genes are distributed among different chromosomes; acrocentric chromosomes in human cells and, in mouse cells, and depending on strain, they can generally be found at chromosomes 12, 16, 18, and 19 (rDNA-chromosomes) and close to centromeric regions ^3^. Due to ribosome biogenesis, the nucleolus structure is influenced by cell states such that highly proliferative cells have larger nucleoli compared to quiescent cells ^6–9^. Accordingly, alterations in nucleolar activities and structure have been well documented under many cellular stress conditions and in several diseases, such as cancer, neurodegenerative disorders, and premature ageing ^10^. For example, nucleolar stress is defined as a condition in which abnormalities in nucleolar structure and function driven by diverse cellular insults, such as nutrient starvation or DNA damage, ultimately leads to activation of stress signalling pathways that downregulate ribosome biogenesis, including rRNA gene transcription, and alter nucleolar structure ^11,12^. However, whether these structural alterations might have an impact on the genome organization around nucleoli remains unexplored, mainly due to technical limitations of current methods to map NADs.

Currently, there are several methods to identify NADs; the biochemical purification of nucleoli followed by sequencing and, more recently, the Nucleolar-DamID that relies on the expression of an engineered nucleolar histone H2B fused to Dam that serves to mark NADs with adenine methylation (H2B-Dam-NoLS) and nucleolar-TSAseq ^13,14^. Although based on different methodologies, these methods revealed that NADs are composed of repressive chromatin domains ^15–19^. However, these methods have also some limitations. First, they cannot be applied to study how genome organization around nucleoli is affected under nucleolar stress ^20,21^. Indeed, under these conditions, nucleoli become frail and cannot be efficiently biochemically purified. On the other side, the consequent downregulation of ribosome biogenesis decreases protein translation, thereby affecting the expression of H2B-Dam-NoLS. Moreover, the physico-chemical alterations of nucleoli upon rDNA downregulation causes the expulsion of the H2B-Dam-NoLS from nucleoli similarly to other nucleolar components ^19^. Finally, these methods have only been applied in cell populations and consequentially they have provided an average nucleolar contact frequency, which might not accurately reflect the structure of the genome around single nucleoli. Thus, so far there is no methodology able to identify NADs under nucleolar stress conditions and in single cells.

To overcome these challenges, we developed NoLMseq, a novel approach that combines laser-capture microdissection (LCM) and DNA sequencing to identify NADs in single cells. LCM is a technique traditionally used for isolating single cells or specific tissue regions^22–26^ that we have expanded it for the organelle-level isolation of single nucleoli. Traditional techniques to study genome organization predominantly rely on either microscopy or DNA sequencing. NoLMseq synergistically integrates strengths of both these approaches, enabling precise examination of the genome architecture surrounding single nucleoli. The application of NoLMseq in mouse embryonic stem cells (ESCs) not only confirmed the general repressive state of NADs but also provided novel insights of genome organization around nucleoli that could not be detected in previous NAD studies since these were based on methods that could only be applied in cell populations. These findings included the detection of different levels of NAD heterogeneity among ESCs that culminates in two distinct cell populations with distinct chromatin states, one more repressed than the other one. NoLMseq also determined that genes detaching from nucleoli during ESC differentiation into neural progenitors are activated and linked to neuronal pathways, implying an active role of the nucleolus in gene repression. Intriguingly, although NADs are generally composed of repressive chromatin domains, NoLMseq detected a high frequency association of the highly expressed ribosomal protein genes with the nucleolus. These findings indicate that the repressive nucleolar compartment can also accommodate transcriptionally active genes and suggest a potential direct role of nucleolar proximity in regulating ribosomal protein genes and, consequently, ribosome biogenesis. Moreover, NoLMseq revealed that NADs prevalently contact nucleoli in a monoallelic manner and that allelic nucleolar contact frequency correlates with gene expression and chromatin states. Finally, we determined how genome structure is impacted under conditions that impair nucleolar integrity, revealing the detachment of a substantial fraction of NADs from nucleoli and their subsequent transcriptional activation. This insight, unattainable with previous technologies, underscores the critical role of nucleolar integrity in genome organization and gene expression, while also illuminating how the genome responds to nucleolar stress.

Together, these results not only provided novel insights of how chromosomes are organized around nucleoli and their remodelling during cell fate commitment and stress-induced nucleolar structural changes but also demonstrated that NoLMseq accurately measures chromosome contacts around single healthy and “stressed” nucleoli. Thus, NoLMseq will be a critical tool to study NADs within biological populations and determine how genome organization responds to nucleolar stress in healthy and disease states.

## Results

### NoLMseq identifies NADs in single cells

To map NADs at a single nucleoli resolution, we combined laser-capture microdissection (LCM) of nucleoli and DNA sequencing. We named this method nucleolus laser microdissection sequencing (NoLMseq). We applied NoLMseq to mouse embryonic stem E14 cells (ESCs), thereby enabling comparisons with NADs from ESC populations, which have recently been identified using Nucleolar-DamID ^19^. ESCs were grown on a thin polyester membrane that can be cut with a fine laser (**Fig. 1a,b**). In brightfield microscopy, the nucleolus appears as a distinct, dark, and dense region within the nucleus, allowing us to visualize and excise it using laser capture microdissection (LCM). The isolated nucleolus was then collected into the cap of a microcentrifuge tube using MMI CapSure technology. The DNA content of each nucleolus was purified and amplified via quasilinear whole-genome amplification (WGA)^27^. To prevent any loss of DNA, WGA was carried out in the same tube in which the microdissected nucleolus was collected. Before proceeding with library preparation and DNA sequencing, the purity of nucleolar DNA was assessed by PCR to determine the presence of rRNA genes, which are located within nucleoli, and the absence of *Tuba1*, an active gene that previous studies have shown to not contact nucleoli^19^ (**Extended Data Fig. 1a**).

**Figure 1.**
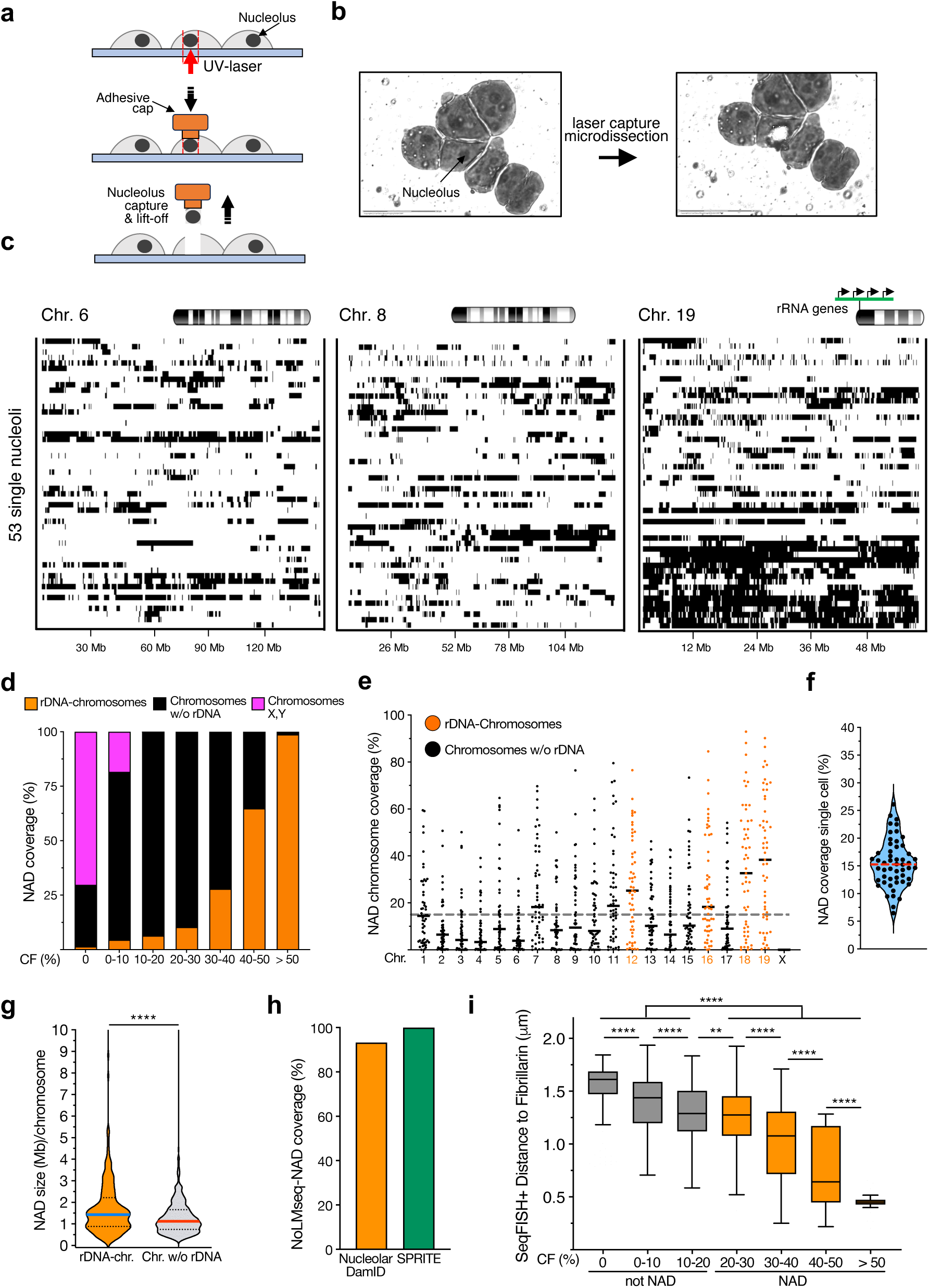
NoLMseq identifies NADs in single cells. **a.** Scheme representing the nucleolar laser capture microdissection (NoLMseq). **b.** Representative images of ESCs before and after nucleolar laser capture microdissection. **c.** Nucleolar contacts maps from 53 single nucleoli for chromosomes 6, 8, and 19. Black bars indicates NADs. **d.** Cumulative histogram of genome-wide nucleolar contact frequency (CF) values with respect to their location at chromosomes containing rRNA genes (rDNA-chromosomes), without (w/o rDNA), and X and Y chromosomes. **e.** Coverage of NADs with CF > 20% on each chromosome in single nucleoli. Dotted grey line indicates the mean NAD coverage in single cells. Black lines indicate the mean on each chromosome. **f.** Violin plot showing NAD coverage in single cells. Red dashed line indicates the median. **g.** Size of NADs located at rDNA-chromosomes and chromosomes not bearing rRNA genes (w/o rDNA). Red and blue lines indicate medians, and the dotted lines indicate quartiles. Statistical significance (*P*-values) was calculated using unpaired t-test (**** < 0.0001). **h.** Coverage of NADs identified by NoLMseq relative to NADs identified in ESC populations with the Nucleolar-DamID ^19^ and SPRITE ^29^ method. **i.** Genome-wide comparison of DNA seqFISH+ distance to exterior of nucleoli marked with Fibrillarin (µm) ^30^ and NoLMseq NAD contact frequency for 3713 paired genomic regions. Tukey boxplot where box limits represent the 25th and 75th percentiles and the black line represents the median. Statistical significance (*P*-values) was calculated using unpaired t-test (** < 0.01; **** < 0.0001).

We generated 53 high-quality single nucleolus contacts maps from ESCs, with a median of 5·10^5^ uniquely mapped reads per nucleolus (**Fig. 1c, Extended Data Figs. 1b and 2**). We binned the genome into 100 kb segments and called NADs using SICER2 ^28^ at a 100kb resolution using genomic DNA as a control. The identified reads over the reference genome showed a bimodal distribution, indicating that loci are either in a “contact” or “no contact” state with nucleoli (**Fig. 1c, Extended Data Fig. 1b**). Since the microdissection of nucleoli will inevitably include genomic sequences above and below nucleoli, we analysed the data by calculating the nucleolar contact frequency (CF) that we defined as the proportion of microdissected nucleoli containing a defined NAD sequence (**Fig. 1d**). We found that NADs from rDNA-chromosomes have higher CF than chromosomes not containing rRNA genes, underscoring the specificity of NoLMseq in identifying NADs. Moreover, chromosome X, which is active in the analysed male ESCs, showed the lowest nucleolar CF. For all downstream analyses, we selected NADs with CF> 20%, a value that corresponds to FDR 0.0009. The analysis of NAD coverage/chromosome of all 53 nucleoli further supported the high propensity of rDNA-chromosomes to contact nucleoli relative to the other chromosomes (**Fig. 1e**). Furthermore, the analysis of 53 microdissected nucleoli appears to be sufficient for robust NAD identification. Calculation of nucleolar contact frequencies and genome coverage using subsets of 20, 30, and 40 randomly selected nucleoli indicates that NAD saturation is reached by 40 nucleoli (**Extended Data Fig. 1c,d**).

The average NAD coverage in single nucleoli was about 15%, much lower than the 30% NAD coverage obtained from previous NAD studies in ESC population ^17,19^ (**Figs. 1e,f**). The data also showed that the average length of NADs of rDNA-chromosomes was 1.85 Mb and significantly higher than the average NAD size from the rest of the chromosomes (1.30 Mb, **Fig. 1g**). The accuracy of NoLMseq was also supported by finding >90% NADs overlapping with previous NAD profiles from ESC populations using Nucleolar-DamID ^19^ and SPRITE method ^29^ (**Fig. 1h**). To further validate the data, we used an orthogonal approach by computing the CF of NADs from NoLMseq genomic data and their distance to Fibrillarin marked nucleoli obtained by seqFISH+ microscopy ^30^ (**Fig. 1i**). We observed that nucleolar CFs measured by NoLMseq were inversely proportional to the distance of NADs to the nucleolar marker Fibrillarin, indicating that NoLMseq accurately measures genomic distance to nucleoli.

To further support the efficacy of NoLMseq, we analysed known NAD features of the identified NADs. It is well known that centromeres tend to be positioned close to the nucleolus in various model systems. Since in mouse cells all chromosomes are acrocentric, we measured the average NAD coverage at centromere-proximal regions at the 5’-quintile portion of all chromosomes and found that it increases with nucleolar CF, further supporting the accuracy of NoLMseq to identify sequences in contact with nucleoli (**Fig. 2a**). Consistent with previous results in ESC populations obtained with Nucleolar-DamID^19^, NADs identified by NoLMseq were significantly depleted of active histone marks relative to sequences within the active A compartment, showed a significant enrichment for the repressive H3K9me2 but not for the other repressive marks H3K27me3 and H3K9me3, and displayed low gene expression levels (**Fig. 2b,c**). Although NADs generally appear to have a repressive chromatin and gene expression state, we could also find cases where highly expressed genes have high nucleolar CFs. In particular, we found high nucleolar CFs for ribosomal protein (RP) genes, which encode proteins for ribosome assembly in nucleoli and are transcribed by RNA Pol II, and tRNA genes, which are transcribed by Pol III (**Fig. 2d**). In contrast, snoRNA genes, which express RNAs playing an important role in rRNA modifications in nucleoli, did not show any preferential location to nucleoli. The nucleolar contacts of RP and tRNA genes was not due to their location at rDNA-chromosomes since most of them were located at chromosomes non containing rRNA genes (92% and 90%, respectively; **Extended Data Fig. 3**). These results raise the possibility that nucleolus proximity could play a role in regulating RP genes and ribosome biogenesis, as well as tRNA, potentially influencing protein translation on a broader scale (**Fig. 5b**). Finally, these data also suggested that not all NADs are necessarily transcriptionally repressed.

**Figure 2.**
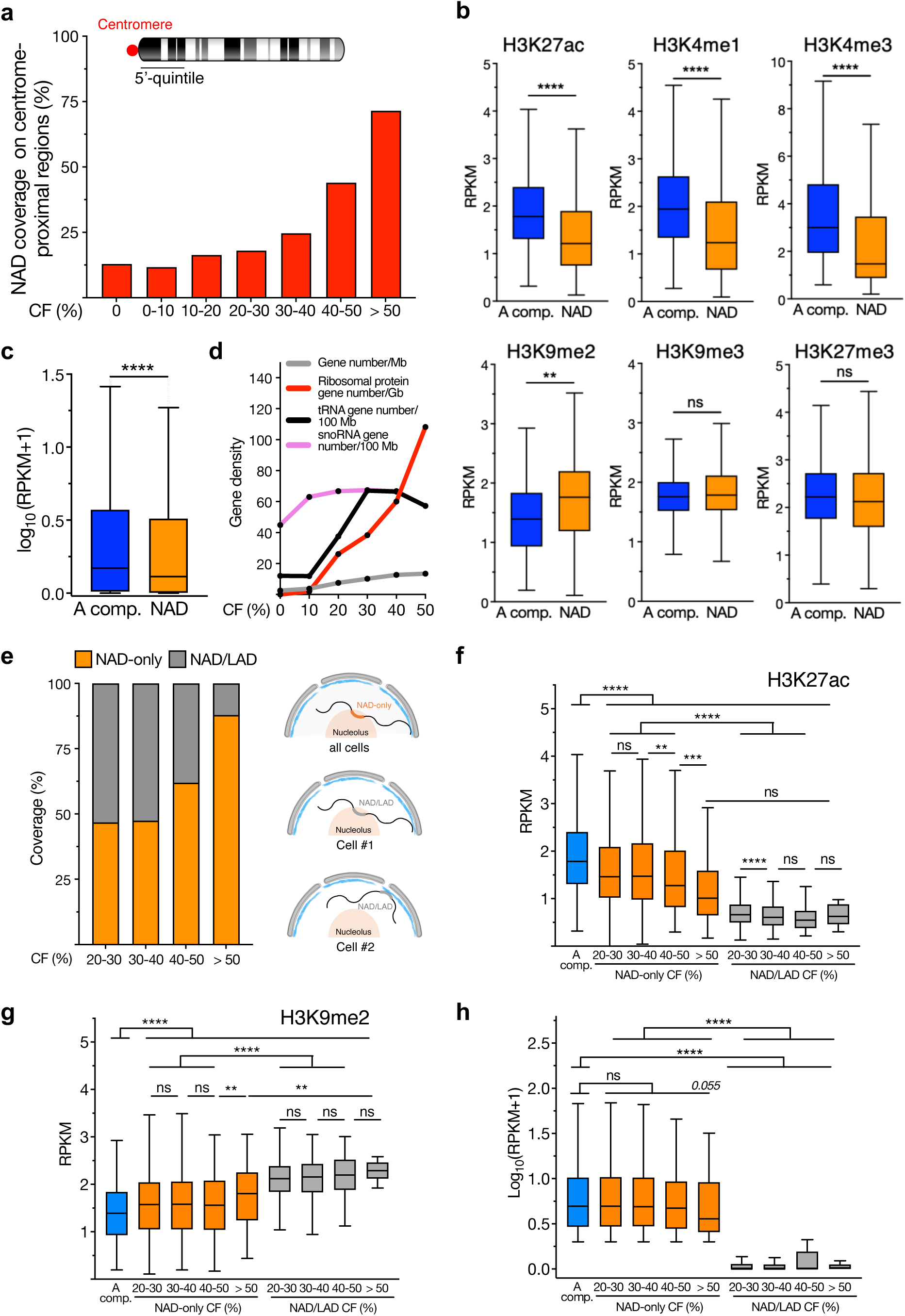
NADs identified by NoLMseq correspond to repressive chromatin domains. **a.** NAD coverage on the centromere-proximal regions (5’ -quintile) as a function of nucleolar contact frequency. **b.** Levels of active histone marks (H3K27ac, H3K4me1, H3K4me3) and repressive histone marks (H3K9me2, H3K9me3, H3K27me3) at NADs relative to genomic regions located at the active A compartment (A comp.). Values are shown as average RPKM. Tukey boxplot where box limits represent the 25th and 75th percentiles. The horizontal line within the boxes represents the median. Statistical significance (*P*-values) was calculated using the unpaired two-tailed t test (**<0.01, ****<0.0001, ns: nonsignificant). **c.** Expression values (RPKM) of genes within the active A compartment (A Comp.) and NADs. Tukey boxplot where box limits represent the 25th and 75th percentiles. The horizontal line within the boxes represents the median. Statistical significance (*P*-values) was calculated using the unpaired two-tailed t test(****<0.0001). **d.** Gene densities of ribosomal protein genes (red), tRNA genes (black), snoRNA genes (pink) and all genes (grey) as a function of nuclear CF. **e.** Left panel shows nucleolar contact frequency for NAD-only and NAD/LAD sequences. Right panel depicts how NAD-only and NAD/LAD contact nucleoli in single cells. **f,g,h.** Levels of H3K27ac (**f**), H3K9me2 (**g**), and gene expression (**h**) at NAD-only and NAD/LAD as a function of nucleolar CF relative to the active A compartment (A comp.). at the active A compartment (A Comp.) and at NAD-only and NAD/LAD relative to nucleolar CF. Values are shown as average RPKM. Tukey boxplot where box limits represent the 25th and 75th percentiles. The horizontal line within the boxes represents the median. Statistical significance (*P*-values) was calculated using the unpaired two-tailed t test (**<0.01, ***< 0.001, ****<0.0001, ns: nonsignificant).

NADs have been previously characterized in two categories; NAD-only (or Type-1) which correspond to sequences contacting nucleoli but not the nuclear lamina (NL) and NAD/LAD (or Type-2) which are regions that can be located both at nucleoli and NL ^13,17–19^. Since this classification of NADs was obtained from ESC population, it has remained elusive whether NAD/LAD regions contact both the nucleoli as well as the NL in the same or different cells. The analysis of single nucleoli data revealed that NAD-only sequences have higher CF values than NAD/LAD, suggesting that NAD/LAD domains can contact the nucleolus in one cell and the NL in another cell, but rarely both nucleolus and NL in the same cell (**Fig. 2e**). We also observed an inverse correlation of NAD-only CF values and H3K27ac content, indicating that strong NADs generally exhibit low levels of active chromatin (**Fig. 2f**). Similarly, NAD-only domains with high nucleolar CF have high H3K9me2 levels (**Fig. 2g**). However, while NAD/LAD regions showed significantly lower gene expression levels relative to genes within the active A compartment, the expression of genes within NAD-only regions was not significantly reduced, albeit there was a tendency to be low expressed when located at regions with the highest nucleolar CF (**Fig. 2h**). These results are consistent with the data described above showing that some active genes can also contact nucleoli.

### ESCs are composed of two populations with distinct nucleolar contacts

NADs obtained from single nucleoli allowed to study the degree of cell heterogeneity in genome organization. We calculated the NAD coverage for each chromosome in single cells and performed hierarchical clustering (**Fig. 3a**). We found two main ESC populations that differ according to the level of nucleolar contacts among chromosomes. The major differences between these two groups were on the level of rDNA-chromosome contacts with the nucleolus, which showed a mutually exclusive relationship. rDNA-chromosomes 12 and 19 had high levels of nucleolar contacts in population 1 whereas they were almost absent in nucleoli of population 2 (**Fig. 3b**). In contrast, rDNA-chromosomes 16 and 18 displayed higher NAD coverage in population 2 than in population 1. In the case of chromosomes not bearing rRNA genes, chromosomes 7, 11, 15, and 2 showed more nucleolar contacts in population 2 than in population 1 whereas chromosomes 3 and 5 preferentially contact nucleoli of population 1. Although ESCs are predominantly in the S phase, the potential influence of cell cycle factors cannot be entirely ruled out. Nonetheless, it is notable that the analysis of histone modifications revealed that NADs enriched in population 1 were significantly depleted of active histone marks and enriched in H3K9me2 relative to NADs of population 2, suggesting different chromatin states between these two ESC populations (**Fig. 3d**). Thus, the data revealed heterogeneity in the ESC genome structure and chromatin configuration that depend on the genome organization around nucleoli.

**Figure 3.**
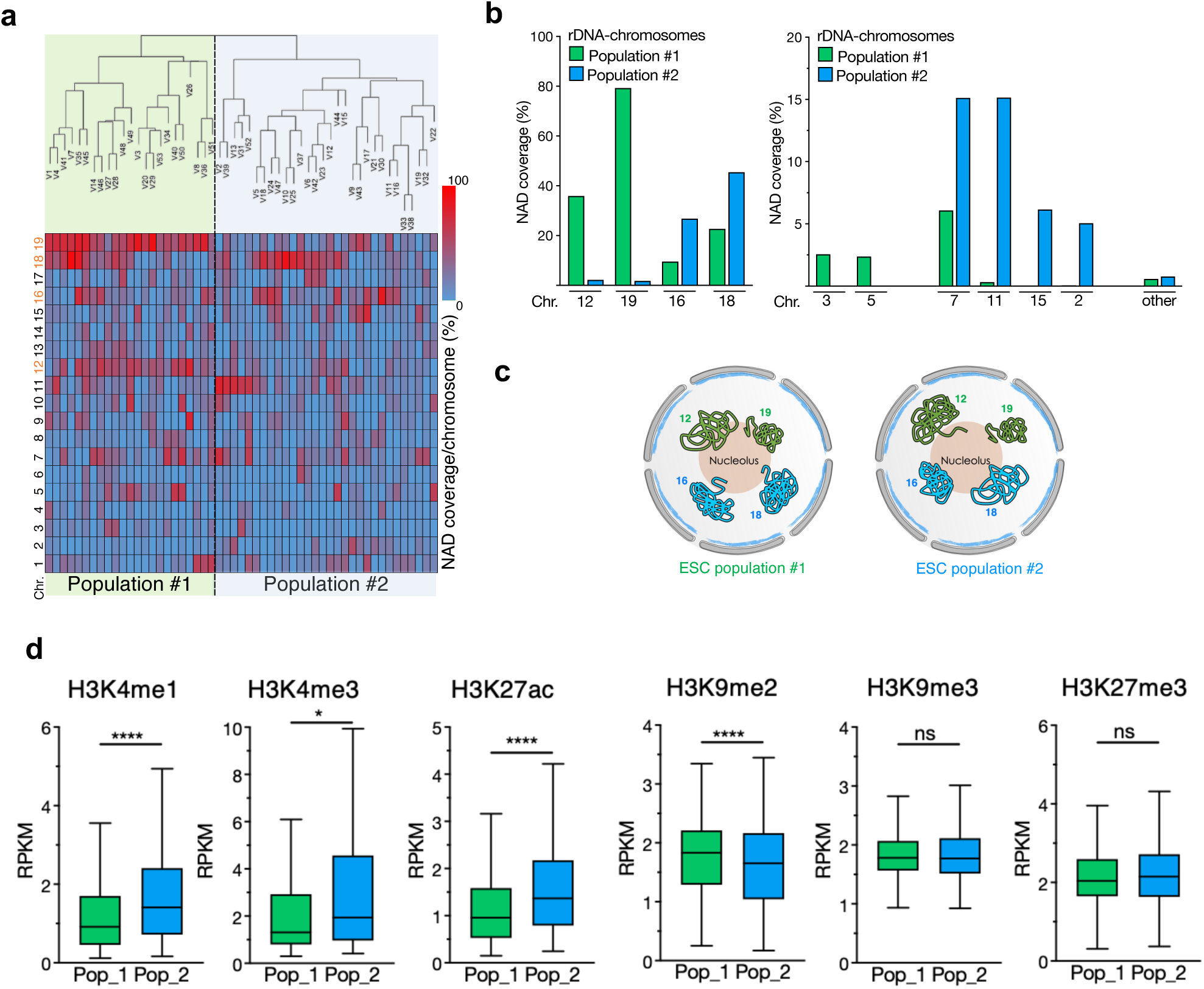
ESCs showed two populations with distinct nucleolar contacts. **a.** Hierarchical clustering of nucleoli according to their NAD coverage/chromosome **b.** NAD coverage at the indicated chromosomes in the top quartile NADs of the two ESC populations. **c.** Scheme representing the contact with nucleoli of the rDNA-chromosomes with the nucleoli in the two ESC populations. **d.** Levels of active histone marks (H3K4me1, H3K4me3, H3K27ac) and repressive histone marks (H3K9me2, H3K9me3, H3K27me3) at NADs enriched in ESC population (Pop) 1 or 2. Values are shown as average RPKM. Tukey boxplot where box limits represent the 25th and 75th percentiles. The horizontal line within the boxes represents the median. Statistical significance (*P*-values) was calculated using the unpaired two-tailed t test (*<0.05, ****<0.0001, ns: nonsignificant).

### Detachment of NADs upon differentiation into NPCs results in gene activation

Next, we asked whether the composition of genomic contacts in single nucleoli could change upon differentiation of ESCs into neural progenitors (NPCs). As in the case of ESCs, in NPCs NADs from rDNA-chromosomes consistently exhibited higher nucleolar CF than regions from other chromosomes (**Fig. 4a**). However, NPCs displayed fewer NADs than ESCs, with a large proportion (75%) being the same NADs as found in ESCs (**Fig. 4b-d**). Moreover, we observed more variability in nucleolus association in ESCs than in NPCs as indicated by the Yule’s Q coefficient, which provides a measure of similarity between single-cell NADs (**Fig. 4e**). The NAD detection in single NPC nucleoli obtained with NoLMseq is consistent with previous studies performed in NPC populations showing that the architecture of chromosomes surrounding the nucleolus is in a more compact and rigid form whereas in ESCs it appears more flexible and variable ^19^. Moreover, these data are consistent with previous reports indicating that ESCs harbour a more open and dynamic chromatin than differentiated cells ^31,32^. Next, we investigated whether genes contacting nucleoli in ESCs but detaching from nucleoli in NPCs (ESC specific NADs, ESC_sp_-NADs) could change their expression state in NPCs. We defined ESC_sp_-NADs those NADs with nucleolar CF > 25% in ESCs and < 10% in NPCs. RNAseq data analysis revealed that ESC_sp_-NADs are less expressed in ESCs than in NPCs, suggesting that the detachment from nucleoli facilitates gene activation (**Fig. 4f,g**). Moreover, the absoluted fold change in gene expression in NPCs compared to ESCs was higher for the upregulated genes, with a large fraction (60%) not expressed in ESCs (**Fig. 4h,i**). Accordingly, GO terms for these upregulated genes were significantly linked to developmental processes, in particularly nervous system development, whereas downregulated genes were mainly implicated in metabolic processes (**Extended Data Fig. 4**). These results ishow that the detachment of genes from nucleoli during ESC differentiation unlocks them for activation, indicating a regulatory role of the nucleolus for the establishment or maintenance of repressive chromatin states.

**Figure 4.**
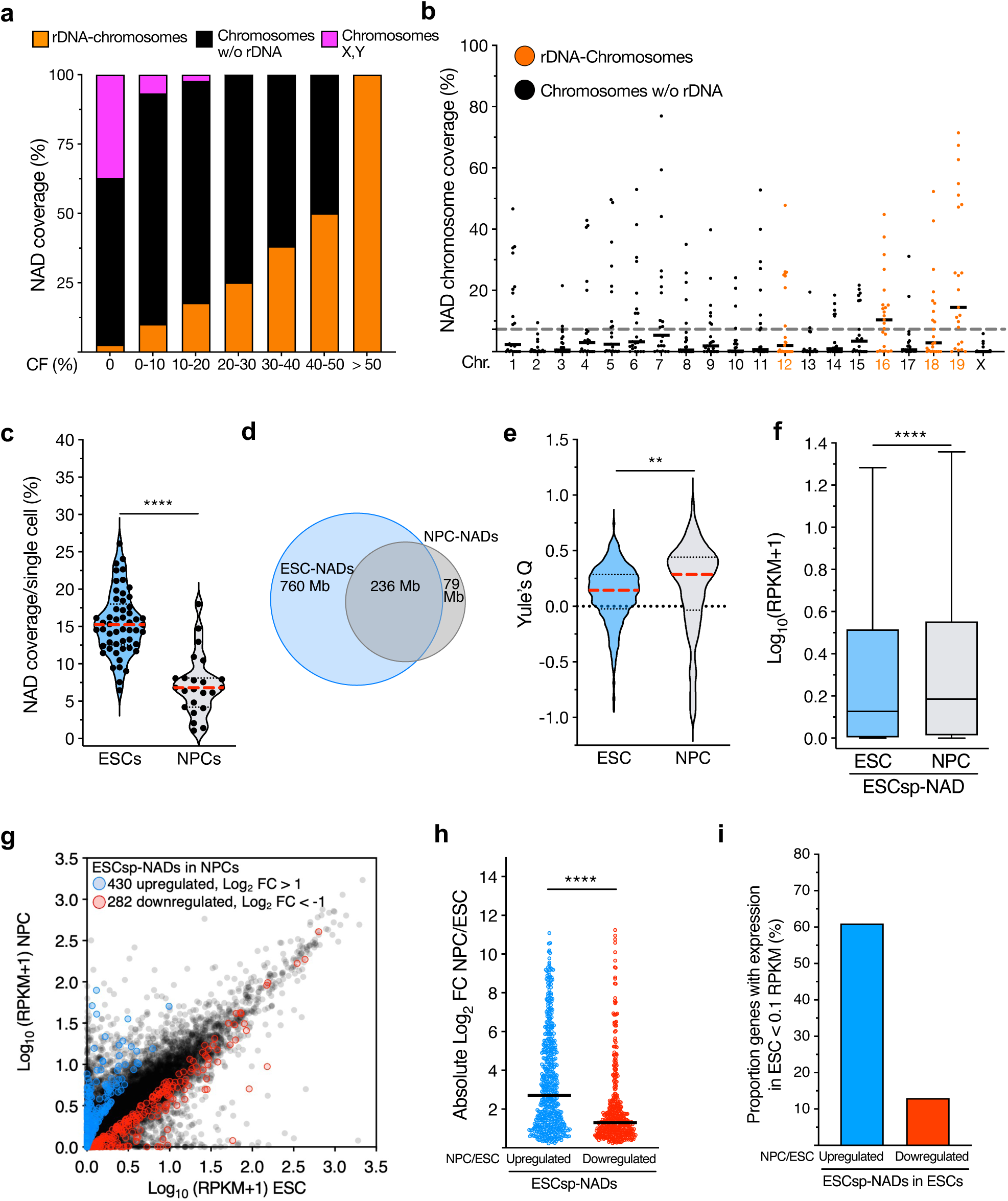
NADs from single nucleoli of neural progenitors. **a.** Cumulative histogram of genome-wide of nucleolar contact frequency measured from 23 microdissected nucleoli with respect to their location at chromosomes containing rRNA genes (rDNA-chromosomes), without (w/o rDNA), and X and Y chromosomes in NPCs. **b.** Coverage of NADs with CF > 20% on each chromosome in single nucleoli of NPCs. Dotted grey line indicates the average NAD coverage in single cells of NPCs. Black lines indicate the mean on each chromosome. **c.** Violin plots showing NAD coverage in single ESCs and NPCs. Data from ESCs were also shown in Figure 1f. Red dashed lines indicate the median and black dotted lines indicate the quartiles. **d.** Venn diagram showing the distribution of NADs in ESCs and NPCs. **e.** Violin plots showing the distribution of Yule’s Q values as a measure of NAD cell-cell similarity in ESCs and NPCs. Red dashed lines indicate the median and black dotted lines indicate the quartiles. **f.** Expression levels of genes located at ESC_sp_-NADs in ESCs and NPCs. Tukey boxplot where box limits represent the 25th and 75th percentiles. The horizontal line within the boxes represents the median. Statistical significance (*P*-values) was calculated using the paired two-tailed t test (****<0.0001). **g.** Scatter plot showing the expression levels of genes at ESC_sp_-NADs in ESCs and NPCs. Expression of genes significantly (P < 0.05) upregulated (blue) and downregulated (red) are shown. **h.** Plot showing the absolute log_2_ fold changes of upregulated and downregulated genes at ESC_sp_-NADs in NPCs. Black lines indicate the median. Statistical significance (*P*-values) was calculated using the unpaired two-tailed t test (****<0.0001). **i.** Proportion of low expressing genes (< 0.1 RPKM) in ESCs that are located at ESC_sp_-NADs and are up- or downregulated in NPCs.

### Nucleolar integrity is required for genomic contacts with nucleoli

Nucleolus structure is highly dynamic and promptly responds to external stimuli that affect ribosome biogenesis and nucleolar structure. For example, downregulation of rDNA transcription by treatment with low doses of Actinomycin D (ActD) induces a spatial reorganization of the nucleolar structure, including the migration of rRNA genes to the nucleolar periphery, forming the so-called nucleolar caps, and the release of the granular component (GC) protein NPM1 from nucleoli (**Fig. 5a**) ^33^. However, until now, it could not be investigated whether these structural changes can impact the organization of the genome around nucleoli mainly due to the technical limitations of the current technologies to map NADs under conditions in which nucleolar integrity is affected. Since nucleoli in ESCs treated with ActD (ESCs+ActD) are still detectable with brightfield microscopy, we reasoned that NoLMseq could be applied to determine how nucleolus integrity affects chromosome contacts with nucleoli. Compared to untreated cells, the average NAD coverage in ESC+ActD significantly decreased from 15% to 10% (**Fig. 5b,c, Extended Data Fig. 5a**). Although NoLMseq revealed a substantial loss of nucleolar contacts across all chromosomes, the majority of NADs retained upon ActD treatment were from the rDNA-chromosomes (**Fig. 5d**). We validated this loss of contacts with nucleoli in ESCs+ActD by quantitative DNA-FISH, using probes targeting four NADs at chromosomes 1, 2, 5, and 19 that were previously identified and validated by Nucleolar-DamID ^19^ (**Fig. 5e,f, Extended Data Fig. 5b**). These are strong NADs since they contact nucleoli in 70-95% of ESCs (**Fig. 5f**). The DNA-FISH probe for the rDNA-chromosome 19 targeted a NAD sequence proximal to the rDNA locus. Accordingly, this NAD remained anchored to nucleoli upon ActD treatment, an expected result since rRNA genes remain close to nucleoli by forming nucleolar caps (**Fig. 5f**). In contrast, DNA-FISH probes targeting NADs at chromosomes 1, 2, and 5 showed a significant detachment of NADs from the nucleoli upon ActD treatment, indicating that nucleolar integrity is crucial for NAD association with nucleoli and supporting the results obtained with NoLMseq. Because of the retainment of rDNA-chromosomes to nucleoli in ESC+ActD, we reasoned that regions from rDNA-chromosomes still able to contact nucleoli in ESC+ActD should correspond to centromere-proximal sequences, which are linearly located close to rRNA genes. Accordingly, NAD coverage of centromere-proximal sequences of rDNA-chromosomes was higher than the corresponding NAD coverage in untreated ESCs, suggesting a reorganization of rDNA-chromosomes structure around nucleoli (**Fig. 5g**). In contrast, NAD coverage of centromere-proximal sequences of chromosomes not bearing rDNA repeats showed a significant reduction compared to NADs of untreated cells, further indicating that the anchoring of these sequences to nucleoli depends on nucleoli integrity. All these results indicate that nucleolar integrity is an important determinant for chromosome organization around nucleoli. To assess whether the detachment of NADs from nucleoli upon nucleolar stress affects gene expression, we performed quantitative RNAseq spiked-in with *ERCC* control RNAs (**Fig. 5h**). Treatment with ActD significantly altered the expression of 5382 genes (log_2_ fold change ≥ 1; *P* < 0.05), of which 2263 genes were upregulated and 3118 were downregulated compared to control cells. Notably, 40% of the upregulated genes corresponded to those detaching from the nucleoli, while only 23% of the downregulated genes were NAD genes that lost contact with nucleoli upon ActD treatment (**Fig. 5i**). The upregulated NAD-genes were linked to signaling pathways involving calcium, PI3K-Akt, and p53, the latter being well known to be activated by RNA Pol I inhibition upon ActD treatment ^34,35^ (**Extended Data Fig. 5c**). These results indicate that the detachment of genes from nucleoli, when nucleolar integrity is compromised, leads to their transcriptional activation (**Fig. 5j**). Together with the reactivation of NAD genes during ESC differentiation into NPCs upon loss of nucleolar contact, these results demonstrate an active role of the nucleolus in repressing gene expression. Moreover, the data demonstrate that NoLMseq is a unique and only genomic method for studying the role of nucleolus in genome organization under nucleolar stress conditions that affect nucleolar integrity.

**Figure 5.**
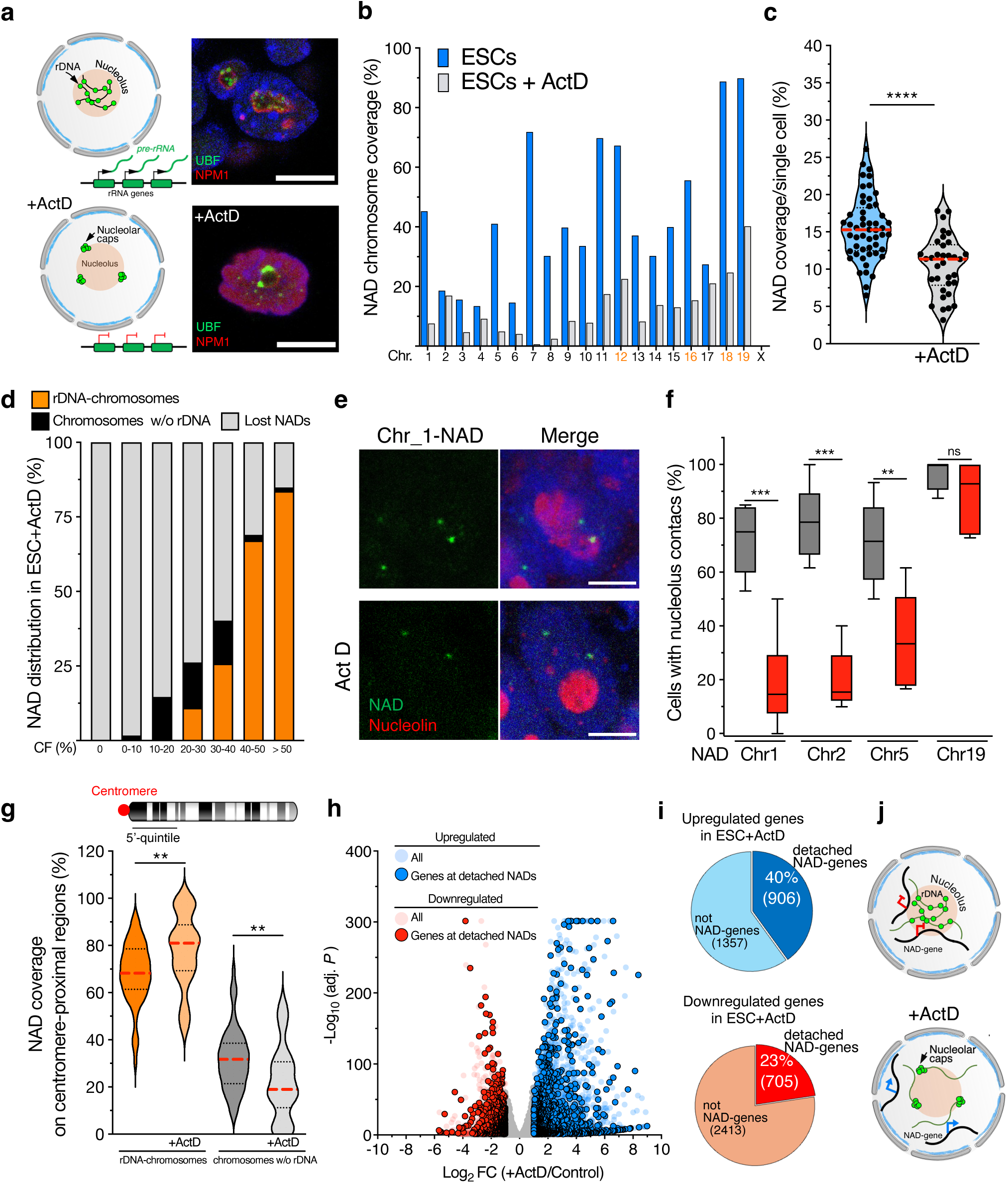
Nucleolar integrity is required for genomic contacts with nucleoli. **a.** Left panel represents alterations in nucleolus structure upon treatment with Actinomycin D (ActD), showing the re-organization of rRNA genes, which form nucleolar caps. Right panel shows immunofluorescence images of the nucleolar markers UBF and NPM1. Scale bar is 9 μm. **b.** Histogram showing the average NAD coverage/chromosome from single nucleoli of untreated ESCs (blue) and 34 microdissected nucleoli from ESCs treated with ActD (grey). **c.** Violin plots showing NAD coverage in single untreated ESCs (blue) and ESCs treated with ActD (grey). Data from ESCs were also shown in Figure 1F. Red dashed lines indicate the median and black dotted lines indicate the quartiles. Statistical significance (P-value) was calculated using the unpaired two-tailed t test (****<0.0001). **d.** Distribution of NADs in ESCs treated with ActD as a function of nucleolar CF across rDNA-chromosomes, chromosomes without (w/o) rDNA, and NADs lost relative to untreated ESCs. **e.** Representative immuno-FISH images for NAD at chromosome 2. Nucleolin serves as nucleolar marker. Scale bar is 5 μm. **f.** Box plots represent the quantifications of immuno-DNA-FISH analyses showing the percentage of cells displaying NADs contacting nucleoli. Error bars represent s.d. Statistical significance (*P*-values) from three independent experiments was calculated using Mann-Whitney test (** < 0.01; *** < 0.001; ns: non-significant). **g.** NAD coverage at the centromere-proximal regions (the first 5’-quintile) between rDNA-chromosomes and chromosomes without (w/o) rDNA. rDNA on untreated ESCs and ESCs treated with ActD. Red dashed lines indicate the median and black dotted lines indicate the quartiles. Statistical significance (P-value) was calculated using the unpaired two-tailed t test (**<0.01). **h.** Volcano plot showing fold change (log_2_ values) in transcript levels of ESC+ActD and control ESCs. Gene expression values of three replicates were averaged and selected for log2 fold changes ≥ 1and adjusted (adj.) *P* < 0.05. Upregulated and downregulated genes at detached NADs are shown. **i.** Pie chart showing the proportion of upregulated and downregulated NAD-genes detached from nucleoli in ESC+ActD relative to all significantly regulated genes (log2 fold changes ≥ 1 and adj. *P* < 0.05. **j.** Schematic representation of alterations in genomic contacts with nucleoli and gene expression following ActD treatment. The upregulation of genes detaching from nucleoli, the formation of nucleolar caps, and the increased frequency of centromere-proximal contacts with rDNA chromosomes are shown.

### Genomic contacts with nucleoli are predominantly mono-allelic

Recent data have started to detect allelic asymmetry in gene expression and 3D-genome organization ^36,37^. So far, however, all studies on NADs were performed using cells whose maternal and paternal chromosomes could not be distinguished from each other, as in the case of the E14 ESC line that derives from a fully inbred homozygous mouse strain. Consequently, the information on NAD coverage has been based on the assumption that both alleles contact nucleoli. To determine the allelic distribution of NADs in single nucleoli, we applied NoLMseq to the male mouse F123-ESC line that was derived from the F1 generation of two fully inbred homozygous mouse strains: *Mus musculus castaneus* (CAST, paternal chromosomes) and *Mus musculus domesticus* S129S4/SvJae (S129, maternal chromosomes)^38^. The F123 DNA sequence has been well characterized and contains high density of single nucleotide variants (SNVs), with an average of 1 SNV every 124 nucleotides across autosomes ^39^. We analysed NAD profiles from 50 microdissected nucleoli of F123-ESCs by aligning them against the F123 genome to identify allele-specific NADs (i.e., NAD_S129_ and NAD_CAST_) (**Fig. 6a**). The vast majority (93%) of total NADs identified in F123-ESCs corresponded to NADs identified in E14-ESCs, indicating that genomic contacts with nucleoli are generally conserved between cell lines with the same developmental stage but from different strains (**Extended Data Fig. 6a**). NADs of F123-ESCs showed a slightly lower genome coverage (11.3%, calculated to haploid genome) relative to 15% NAD coverage measured in E14-ESCs (**Fig. 6B**). Surprisingly, we found that a large fraction of NADs (86%) are mono-allelic (**Fig. 6a-c**). Monoallelic NADs were equally distributed among S129 and CAST chromosomes (**Extended Data Fig. 6b**). The distinction between monoallelic and biallelic NADs was also evident from nucleolus CF analysis, which showed that biallelic NADs have a higher CF than monoallelic NADs whereas NAD-only and NAD/LAD regions display similar content (**Extended Data Fig. 6c**). We validated these results by DNA-FISH for two strong NADs, one located at chromosome 5 and another located at the rDNA-chromosome 19 and close to the rDNA locus (**Fig. 6d**). The NAD of chromosome 5 showed monoallelic contacts in almost all the cells. Remarkably, even the NAD of chromosome 19 located close to the rDNA locus showed mono-allelic contacts with nucleoli in 50% of the cells, supporting the results of NoLMseq. This prevalent mono-allelic NAD distribution resulted in a much lower NAD coverage per cell (5.6%, 297 Mbp) compared to the previously assumed bi-allelic distribution (**Fig. 6e**). To determine whether mono- and bi-allelic NADs reflect differences in chromatin modifications, we performed ChIPseq analysis for H3K9me2, H3K27ac, and H3K4me3 in F123-ESCs (**Fig. 6f**). We found that bi-allelic NADs are in a more repressed chromatin states than mono-allelic NADs since they display reduced levels of H3K27ac and H3K4me3 and higher levels of H3K9me2, the latter the repressive histone mark characterizing NADs in ESCs ^19^. Similarly, RNAseq analyses showed a significantly lower expression of genes located at bi-allelic NADs (**Fig. 6g**). Among genes located at bi-allelic NADs, we found cell type specific transcription factors, such as *Gata6* and *TAF4b*, as well as genes linked to metabolic pathways and neuroactive ligand-receptor interactions (**Extended Data Fig. 6c**). These results further indicate that high-frequency contact with nucleoli, even at the allelic levels, results in a more repressive state.

**Figure 6.**
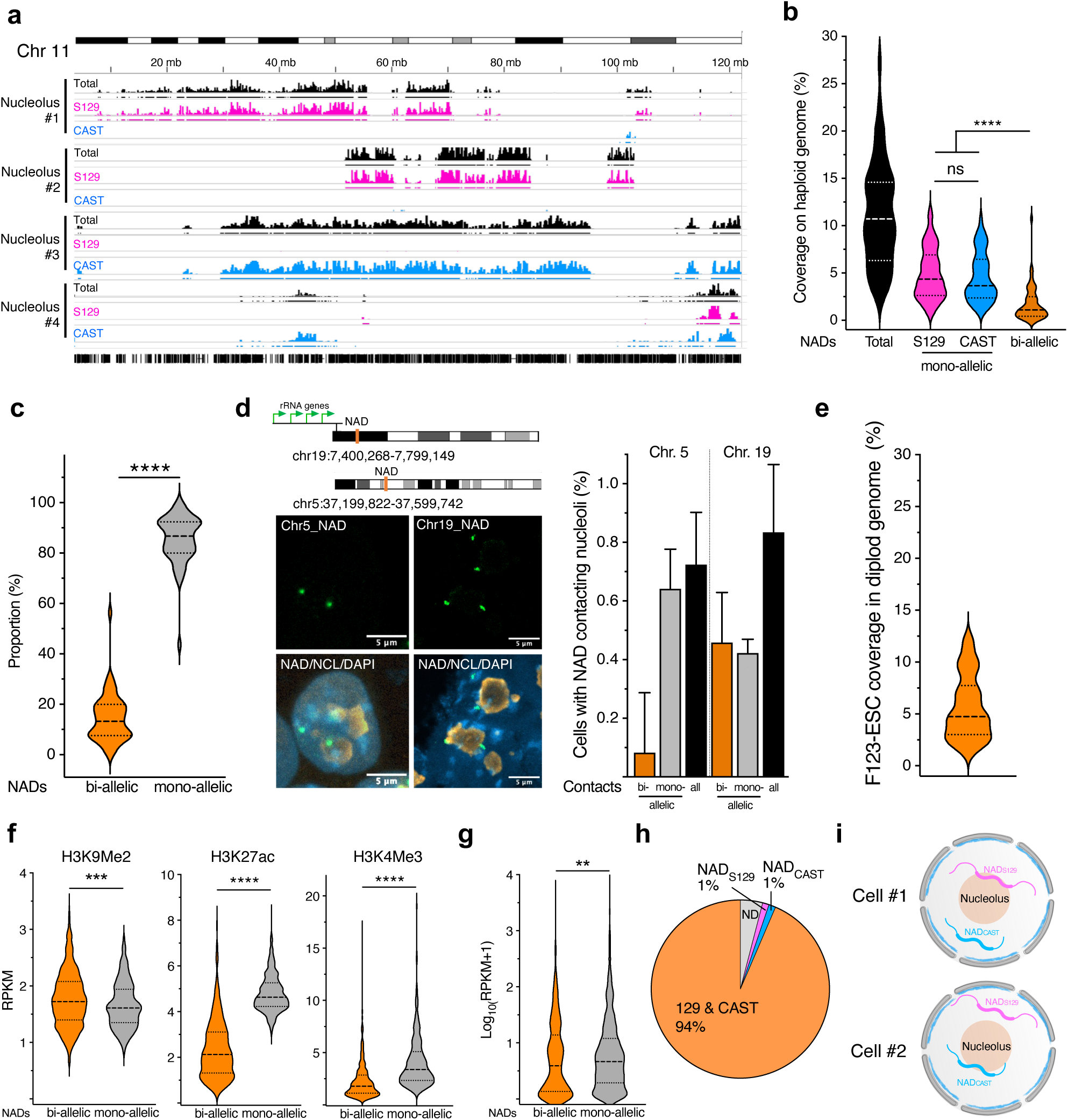
Genomic contacts with nucleoli are mono-allelic. **a.** NoLMseq tracks displaying NAD sequences from four microdissected single nucleoli from hybrid F123-ESCs (S129/CAST) on the total mm10 genome (black), maternal S129 genome (pink) and paternal CAST genome (blue). Bars below the tracks indicate NADs called on respective genomes. **b.** NAD coverage from single nucleoli calculated over the haploid genome. Median and quartiles are shown. Statistical significance (P-value) was calculated using the unpaired two-tailed t test (****<0.0001, ns: non-significant). **c.** Proportion of bi-allelic and mono-allelic NADs in single nucleoli. Black dashed lines indicate median and black dotted lines indicate quartiles. Statistical significance (P-value) was calculated using the unpaired two-tailed t test (****<0.0001). **d.** Left panel shows DNA-FISH images of a NAD located at chromosome 19. Right panel is the quantification of bi-allelic and mono-allelic contacts with nucleoli. Scale bar is 5 μm. Error bars indicate s.d. **e.** NAD coverage from single nucleoli over the diploid genome. Black dashed line indicates the median and black dotted lines indicate the quartiles. **f,g.** Levels of H3K9me2, H3K27ac, and H3K4me3 (**F**) and gene expression (**G**) at bi-allelic and mono-allelic NADs. Values are shown as average RPKM. Violin plots where black dashed lines indicate the median and black dotted lines indicate the quartiles. Statistical significance (*P*-values) was calculated using the unpaired two-tailed t test (**<0.01; ***<0.001; ****<0.0001). **h.** Pie chart showing the amounts of paternal (NAD_CAST_) and maternal (NAD_S129_) NADs. **i.** Model showing that mono-allelic distribution of NADs at nucleoli of single cells is generally not parent specific.

Next, we asked whether the mono-allelic distribution of NADs might be parent specific. We applied stringent criteria to assign parent specific NADs, considering, for example, that a specific NAD_S129_ should have >25% nucleolar CF with the S129 genome and <10% nucleolar CF with the CAST genome (**Fig. 6h**). However, very few NADs (1% on both alleles) showed a parent specific distribution. This indicates that, generally, in one cell it is the paternal allele contacting nucleoli while in another cell it is the maternal allele (**Fig. 6h**).

## Discussion

We described NoLMseq, a method able to measure genome organization around nucleoli in single cells. LCM has been widely used in various omics approaches for isolating single cells or specific tissue regions^22–26^. NoLMseq has adapted this technology to study subcellular genomic compartments, specifically the nucleolus, enabling precise examination of the genome architecture surrounding the nucleolus in single cells.

Relative to previous NAD methods, NoLMseq not only detects NADs in single nucleoli but it can also be applied to study genome organization around nucleoli under conditions of nucleolar stress, which profoundly alter nucleolar structure ^6–9^ and that could not be examined with current methods, making elusive how genome structure around nucleoli responds to nucleolar stress. The data revealed a profound reorganization of the genome around “stressed” nucleoli, such an increase in nucleolar contacts of centromere-proximal sequences of rDNA-chromosomes while the rest of the chromosomes lose their contact. The genomic data obtained by NoLMseq are also consistent with recent image analyses showing that perturbation of nucleolus structure upon downregulation of the RNA polymerase I subunit RPA194 reduced centromere-nucleolus interactions ^40^. These results indicate that nucleolar structure and integrity plays an important role in genome organization and future studies will address alterations in chromosomal interactions with nucleoli under pathological stress conditions impacting ribosome biogenesis and how these perturbations are caused and their functional significance. The results also showed that NADs are very dynamic and NoLMseq represents an unique technique able to capture NADs at the present state rather than averages of the past and present states as in the case of DamID as recently discussed ^41^.

The detection of NADs in single nucleoli using NoLMseq has proven to be highly precise compared to previous methods and capable of providing unexplored insights in genome organization. For example, NoLMseq determined that active genes, such as ribosomal protein genes and tRNA genes, also have high frequency contacts with nucleoli. Interestingly, previous imaging analyses in living or fixed yeast cells reported the proximity of some ribosomal protein genes and tRNA genes to nucleoli ^42,43^, suggesting that the nucleolus could act as a compartment for the regulation of the expression of these key components of the translation machinery. The presence of some active genes at nucleoli is also consistent with a recent study showing that the localization of genes at the repressive B compartment or proximal to nucleoli does not necessarily preclude transcription ^44^.

The accuracy of NoLMseq in detecting NADs relative to cell population studies is also shown by the identification of ESC_sp_-NADs that contain genes activated in NPCs, which could not be detected in previous cell population studies ^19^. Similarly, the results showing the activation of genes upon losing contact with nucleoli upon ActD treatment indicate that the nucleolus serves an active role as a repressive compartment. Because of the single nucleolus detection, NoLMseq revealed that ESCs display a certain heterogeneity in genomic contacts, with nucleoli showing two major cell populations with distinct chromatin features. Whether the presence of such populations, one more repressed than the other one, might reflect differences in gene expression, pluripotency states, or other functions remains to be investigated. These results are also consistent with recent data showing heterogeneity of nuclear organization between individual single ESCs ^45^. Finally, the application of NoLMseq in the hybrid F123-ESC line showed that genomic contacts with nucleoli are mainly monoallelic. This result clearly provided a more accurate value of NAD coverage, which was previously calculated based on either a haploid genome or by assuming that all NADs are bi-allelic in the diploid genome. The average NAD coverage 5.6% (i.e., ca. 297 Mbp) per cell obtained in the F123-ESC line is much lower than the 30% NAD coverage measured in ESC population studies using both purified nucleoli and Nucleolar-DamID methods ^17,19^. Although this calculated NAD coverage value does not include repetitive sequences such as major and minor satellites, which are known to contact nucleoli but are not present in the reference genome, it clearly represents a more realistic scenario than to have one third of chromosomes contacting nucleoli. Moreover, the data showed that bi-allelic NADs have high nucleolar CF are in a more repressive state than monoallelic NADs, further supporting the efficacy of NoLMseq in identifying NADs and the nucleolus acting as a repressive compartment.

Together, the results demonstrate that NoLMseq not only accurately measures chromosome contacts around single nucleoli but also provides novel insights into genome organization around nucleoli that could not be detected by previous studies. We predict that NoLMseq will not only be a critical tool to study genome organization around the nucleolus in biological populations but also to study how the 3D-genome organization responds to nucleolar stress in disease states and understand the functional consequence of these alterations.

## Materials and Methods

### Cell lines

129 mouse embryonic stem cells (E14 line) were cultured in serum medium containing Dulbecco’s modified Eagle’s medium + Glutamax (Life Technologies), 15% FCS (Life Technologies; Cat no. 10270106 FBS South American), 1× MEM NEAA (Life Technologies), 100 μM β-mercaptoethanol, recombinant leukemia inhibitory factor, LIF (Polygene, 1000 U/ml), 1× penicillin/streptomycin (Life Technologies). ESCs were seeded at a density of 50,000 cells/cm2 in culture dishes (Corning CellBIND surface) coated with 0.1% gelatin without a feeder layer. Propagation of cells was carried out every 2 days using enzymatic cell dissociation. ESCs were treated with Actinomycin D (10ng/ml) for 24h before downstream analyses.

Neural progenitor cells were generated from ESCs, according to a previously established protocol ^46^. In brief, differentiation used a suspension-based embryoid bodies formation (Bacteriological Petri Dishes, Bio-one with vents, Greiner). The neural differentiation media (DMEM, 10% fetal calf serum, 1× MEM NEAA, 2 mM Pen/Strep, β-mercaptoethanol, and sodium pyruvate) was filtered through 0.22 μm filters and stored at 4 °C. During the 8-day differentiation procedure, media was exchanged every 2 days. In the last 4 days of differentiation, the media was supplemented with 2 μM retinoic acid to generate neural precursors that are Pax-6-positive radial glial cells.

F123-ESCs (mouse male, hybrid cell line, F1 S129/Jae and Cast mouse cross) ^38^ were cultured on a layer of mitotically inactivated feeder murine embryonic fibroblasts under standard conditions (DMEM, supplemented with 15 % KSR, 1x Glutamax, 10 mM non-essential amino acids, 50 µM beta-mercaptoethanol, 1,000 U/ml LIF). Media was changed every 24h. Before harvesting, mESCs were passaged onto feeder-free ESGRO-gelatin (EMD Millipore SF008). coated plates for at least 2 passages to remove feeder cells. Cells were harvested after approximately 48 h at 70 – 80 % confluency. F123-ESCs were a gift from Bing Ren, University of California, San Diego.

### NoLMseq

For laser-microdissection, cells were grown on ESGRO-gelatin coated and UV-irradiated PET membrane slides (1.4um thick, RNase-free, MMI, 50102) for 24h. Cells were washed twice with 1X PBS and fixed with EM-grade freshly-depolymerised 3% PFA in 1X PBS for 30mins. The slides were then placed in 70% ethanol for 1min and air dried. The slides were then washed with filtered ultra-pure H_2_O (3x 1 min each) and air dried. Single nucleoli were laser microdissected with the MMI Laser Capture Microdissection Nikon microscope (Molecular Machines & Industries) into single 0.2 ml tubes (transparent MMI caps, 50208). Nucleoli were identified by bright-field imaging. A 0.2ml cap filled with transparent adhesive material was lowered onto the cells. A laser was used to cut the PET membrane surrounding each nucleolus and the cut PET membrane section along with the single nucleolus was lifted off by the sticky cap. Each single nucleolus was collected in a separate cap and stored for further processing. Identified nucleoli using bright-field microscopy were initially focused using Köhler illumination to achieve optimal contrast for accurate visualization of the nucleoli. To ensure precision, the stage was adjusted with camera-stage alignment, maintaining an inclination of less than ±0.1 degrees. Paraxial lens offset was employed to align the optical axes of all objectives. Laser alignment was calibrated by firing a single shot, adjusting the pixel center of the imaging system to coincide with the laser beam center. Laser cutting parameters, such as cut velocity, laser focus, and laser power, were then optimized. The cut velocity was set at 20 μm/s, laser power at 0.001% of maximum capacity, and the focus was calibrated for each cut to achieve a cut diameter of 0.25 μm thickness, ensuring reproducibility. Prior to nucleolus excision, the laser was refocused on an empty area of the slide, where a cut similar in size to the nucleolus was drawn. The laser’s inner ring was positioned 0.25 μm from the edge of the drawn shape. When the cap was lowered onto the membrane with the sample, any membrane displacement (typically a few microns) was corrected using the Cap Z offset. To maintain consistent focus during slide movement, slide surface focusing function of the microscope was employed to account for potential topological differences in the membrane. Upon identifying a cell with a distinct nucleolus, a 0.2 mL cap containing a transparent adhesive was lowered onto the sample. The inner ring of the laser used to cut the PET membrane is positioned approximately 0.25 μm from the outer edge of the nucleolus. The excised PET membrane, along with the nucleolus, was lifted off using the adhesive cap. The cap was lowered onto an empty area of the membrane to check for the sample was correctly excised. Any caps with improper cuts or those lacking a clearly identifiable nucleolus were discarded. Only caps containing correctly isolated nucleoli were retained for further processing and analysis. After collection, a new cap was added, and the entire focusing and cutting process was repeated to ensure consistency across samples. Each nucleolus was collected in a separate cap for further processing.

Whole-genome amplification (WGA) was carried out using the MALBAC single cell WGA kit (YK001A) following manufacturer’s recommendations with minor modifications. The entire WGA reaction was carried out in the same 0.2ml tube with a single nucleolus. Another tube, with an empty section was used as a negative control whereas 30 pg of whole cell DNA were used as a positive control for the WGA reaction. The cell lysis reaction was first carried out by using a heated lid at 50 °C for 50 mins and then samples were collected by briefly spinning and proceeding with the cell lysis reaction again at 50 °C for 50 mins, followed by inactivation at 80 °C for 10mins. The reaction is then cooled at 4 °C. The lysis step is followed by a pre-amplification step which involves the quasilinear amplification using random primers at 20 °C progressively with 10 °C steps till 70 °C. After cooling, the samples were subjected to a final amplification step by 20 cycles of PCR.

WGA-amplified DNA was purified using a Qiagen MinElute PCR Purification Kit and eluted in 140 μl ultrapure library-grade H_2_O. The amplified nucleolar DNA was checked on a 2% agarose gel to select positive samples showing smears corresponding to amplified DNA similar to the positive control. The positive samples were further checked for the presence of rDNA and the absence of Tubulin gene by PCR on an agarose gel. The samples which passed this quality control check were then used for library preparation and sequencing. 120 μl of the reaction was first sonicated in split-cap tubes by Covaris E220 to obtain fragments around 300-500nt in length (Peak power- 75, Duty factor- 20, cycle/bursts- 1000, duration- 35s). The samples were then purified and concentrated with AMPure XP beads (Beckman Coulter, A63881) and used for Illumina library using the NEBNext ultra II kit (NEB E7600). Briefly, NoLMseq samples (10-100 ng) were end-repaired and polyadenylated before the ligation of Illumina compatible adapters. The adapters contain the index for multiplexing. Library concentrations were estimated using a Qubit® 4 fluorometer (Thermo Fisher Scientific) and the 4200 TapeStation system and libraries were pooled together in batches averaging 40 samples. Each library pool was sequenced in single end 100bp in one lane of Novaseq6000 to get an average 10 million reads per library.

For data analysis, NoLM-seq reads were aligned to the mouse mm10 reference genome using Bowtie2 (version 2.5.0; ^47^) with default parameters. PCR duplicates were removed using samtools ^48^. Positive NAD bins were called using SICER2 ^49^, using the sequenced gDNA file as the control file and the parameters w=100000, g=100000, fdr=0.0001. Bigwig files were created from the bam files using bamCoverage from deeptools ^50^. For F123-ESCs, N-masked genome was created using the mm10 reference genome and the Variant Call *Format* (VCF) file from the mouse genome project containing the SNP information for all strains. The SNPsplit package ^51^ was used to read the VCF file and extract the SNP information for the S129/SvJae and CAST strains and N-mask for the mm10 genome. The reads were aligned using Bowtie2 to this N-masked reference genome using default parameters. SNPsplit was then used to tag the SNPs for S129/SvJae and CAST alleles in the bam file and sort the haplotype-specific bam files for both the alleles. bamCoverage and Sicer2 positive NAD calling was performed independently on the allele-specific bam files. Integrative Genome Viewer (IGV, version 2.15.4) ^52^ was used to visualize and extract representative single nucleolus NoLMseq bigwig and bed tracks.

Nucleolar contact frequency was calculated at a 100kb resolution. We defined as NADs sequences with a > 20% nucleolar contact frequency that correspond to a false discovery rate (FDR) 0.0009 or 0.09%, meaning that more than 99% of called NADs are truly NADs. This was calculated using the cumulative distribution function (CDF) P(X ≤ k) = ∑ 𝑖 = 0𝑘(𝑖𝑛)𝑝𝑖(1 − 𝑝)𝑛 − 1 where “n” is the number of single nucleoli samples, “p” is the probability that a false-positive region is present in the sample, “k” is the contact frequency threshold. For the hierarchical clustering of NADs, NAD chromosome coverages in single nucleoli were used for clustering single nucleoli based on divisive clustering (Diana) using the “cluster” package (version 2.1.6) in R.

### RNAseq

Total RNA was purified with TRIzol reagent (Life Technologies). The quality of the isolated RNA was determined with a Qubit® 4 fluorometer (Life Technologies, California, USA) and 4200 TapeStation system (Agilent, Santa Clara, California, USA). Only those samples with a 260 nm/280 nm ratio between 1.8–2.1 and a 28S/18S ratio within 1.5-2 were further processed. The TruSeq Stranded mRNA (Illumina, Inc, California, USA) was used in the succeeding steps. Briefly, total RNA samples (100–1000 ng) were polyA enriched and then reverse-transcribed into double-stranded cDNA. The cDNA samples were fragmented, end-repaired and adenylated before ligation of TruSeq adapters containing unique dual indices (UDI) for multiplexing. Fragments containing TruSeq adapters on both ends were selectively enriched with PCR. The quality and quantity of the enriched libraries were validated using Qubit® 4 fluorometer. The product is a smear with an average fragment size of approximately 260 bp. Libraries were normalized to 10 nM in Tris-Cl 10 mM, pH8.5 with 0.1% Tween 20. The NovaSeq X (Illumina, Inc, California, USA) was used for cluster generation and sequencing according to standard protocol. Sequencing was paired end at 2 × 150 bp.

The quality of the reads was checked by FastQC, a quality control tool for high throughput sequence data ^53^. The quality of the reads was increased by applying: a) SortMeRNA ^54^ (version 2.1) tool to filter ribosomal RNA; b) Trimmomatic ^55^ (version 0.40) software package to trim the sorted (a) reads. N-masked genome was created using the mm10 reference genome and the VCF file from the mouse genome project containing the SNP information for all strains. The SNPsplit package ^51^ was used to read in the VCF file and extract the SNP information for the S129/SvJae and CAST strains and N-mask the mm10 genome. The sorted (a), trimmed (b) reads were mapped against the Nmasked mouse genome (mm10) using the default parameters of the STAR (Spliced Transcripts Alignment to a Reference, version 2.6, ^56^). SNPsplit was then used to tag the SNPs for S129/SvJae and CAST alleles in the bam file and sort the haplotype-specific bam files for both the alleles. For each gene, exon coverage was calculated using a custom pipeline and then normalized in reads per kilobase per million (RPKM) ^57^.

Cells were treated with Actinomycin D (10 ng/mL) for 24 hours, after which 1×10⁶ cells were collected for RNA purification using the Zymo RNA extraction kit (Zymo Research). Three biological replicates were prepared for both ActD-treated and untreated conditions. RNA isolation was performed according to the manufacturer’s instructions. All samples were then spiked-in with ERCC control RNAs (Thermo Fisher Scientific), following the manufacturer’s recommendations.

Library preparation was carried out using the TruSeq Stranded mRNA kit (Illumina, Inc., California, USA) as described previously. The absolute abundance of mRNA transcripts was assessed using the ERCC92 RNA spike-in control^58^. Most of the 92 ERCC transcripts showed consistent counts between untreated and ActD-treated samples. Reads were aligned to the mouse genome using STAR and a custom pipeline was used to calculate RPKM values.

### ChIPseq

ChIP analysis was performed as previously described ^59^. Briefly, 1% formaldehyde was added to cultured cells to cross-link proteins to DNA. Isolated nuclei were then lysed with lysis buffer (50 mM Tris-HCl, pH 8.1, 10 mM EDTA, pH 8, 1% SDS, 1X protease inhibitor complete EDTA-free cocktail, Roche). Nuclei were sonicated using a Bioruptor ultrasonic cell disruptor (Diagenode) to shear genomic DNA to an average fragment size of 200 bp. 20 μg of chromatin was diluted to a total volume of 500 μl with ChIP buffer (16.7 mM Tris-HCl, pH 8.1, 167 mM NaCl, 1.2 mM EDTA, 0.01% SDS, 1.1% Triton X-100) and incubated overnight with the ChIP-grade antibodies against H3K9me2, H3K9me3, H3K27me3, H3K4me1, H3K4me3 and H3K27ac. After washing, bound chromatin was eluted with the elution buffer (1% SDS, 100 mM NaHCO_3_). Upon proteinase K digestion (50 °C for 3 h) and reversion of cross-linking (65 °C, overnight), DNA was purified with phenol/chloroform, and ethanol precipitated. The quality and quantity of the isolated DNA were determined with a Qubit® 4 Fluorometer (Life Technologies, California, USA) and the 4200 TapeStation system. Briefly, ChIP samples (1 ng) were end-repaired and polyadenylated before the ligation of Illumina compatible adapters. The adapters contain the index for multiplexing. The quality and quantity of the enriched libraries were validated using Qubit® 4 Fluorometer and the 4200 TapeStation system. The libraries were pooled to 10 nM in Tris-Cl 10 mM, pH8.5 with 0.1% Tween 20. Sequencing was performed on the NovaSeq 6000 for 100bp single end reads (Illumina, Inc, California, USA).

For F123-ESCs, N-masked genome was created using the mm10 reference genome and the Variant Call Format (VCF) file from the mouse genome project containing the SNP information for all strains. The SNPsplit package ^51^ was used to read the VCF file and extract the SNP information for the S129/SvJae and CAST strains and N-mask for the mm10 genome. ChIPseq reads were aligned to the N-masked mm10 reference genome with Bowtie2 (version 2.5.0; (version 2.5.0; ^47^) using default parameters. PCR duplicates were removed using samtools ^48^. SNPsplit was then used to tag the SNPs for S129/SvJae and CAST alleles in the bam file and sort the haplotype-specific bam files for both the alleles. Bigwig files were created from the bam files using bamCoverage from deeptools ^50^ using default parameters. Integrative Genome Viewer (IGV, version 2.15.4) ^52^ was used to visualize and extract representative ChIPseq tracks.

### Gene Ontology (GO) enrichment

GO enrichment analysis of genes was performed with DAVID 6.8 ^60^. All genes in *Mus musculus* were used as the background.

### DNA-FISH

DNA-FISH probes for chromosomes 1, 2, 5, and 19 were generated as previously described ^19^. The probes were obtained with oligopaint libraries that were constructed the PaintSHOP interface created by the Beliveau lab and were ordered from CustomArray/Genscript in the 12 K Oligopool format. Each library contains a universal primer pair used to amplify all the probes in the library, followed by a specific primer pair hooked to the 40-46-mer genomic sequences, for a total probe of around 124-130-mers. Oligopaint libraries were produced by emulsion PCR from the pool, followed by a “two-step PCR” and lambda exonuclease as described before ^19,61^. Specifically, the emulsion PCR with the universal primers allowed amplifying all pool of probes in the library. The “two-step PCR” led to the addition of a tail to the specific probes bound to the Alexa Fluor 488 fluorochrome. All oligonucleotides were purchased from Integrated DNA Technologies (IDT, Leuven, Belgium).

DNA-FISH was adapted from the protocol of Cavalli’s lab ^62^. Cells fixed with 4% PFA for 10 min at room temperature 24h after seeding. Cells were washed three times with 1X PBS and then permeabilized with 0.5%Triton-X100/PBS for 10 min on ice, followed by 10 min at room temperature. After three additional washes with 1X PBS, cells were incubated with 0.1 M HCl for 10 min at room temperature, washed twice with 2X SCCT, and twice with 50% Formamide-2X SCCT for 20min, firstly at room temperature and then at 60°C. Probe mixture contained around 45pmol of Oligopaint probe, 0,8µL of ribonuclease A (ThermoScientific, 10 mg/mL), and FISH hybridization buffer for a total mixture volume of 20uL, which was directly added on the slide. Cell DNA was denaturated at 83 °C for 8 min, and hybridization was performed in a humid dark chamber overnight at 42 °C. Cells were washed with 2X SCCT for 15 min once at 60°C and once at room temepature, once with 0.2X SCC for 10 min, twice with 1X PBS for 2 min, and three times with 1X PBT for 2 min. Slides were then incubated for 1 h at room temperature with blocking solution in a dark humid chamber, and overnight at 4 °C with primary antibody against Nucleolin. Cells were washed four times with 1X PBT at increasing incubation times (1 × 2 min, 1 × 3 min, 2 × 5 min) and then incubated for 2 h at room temperature with goat anti-mouse IgG (H + L) Alexa Fluor 546 in a dark humid chamber. Slides were washed four times with 1X PBT at increasing incubation times (1 × 2 min, 1 × 3 min, 2 × 5 min), three times with 1X PBS for 2 min. Then, slides were mounted with Vectashield DAPI mounting media (Vector, H-1200) and, after drying, stored at 4 °C.

DNA FISH/IF samples were imaged using a Zeiss LSM 980 Airyscan, with a z-stack collected for each channel (step size, 0.3 um), using the oil objective Plan-Apochromat/1.40. Images were processed by ImageJ (version 2.14.0/1.54F). The individual cells were identified by Hoechst/DAPI staining and cells containing signal for DNA-FISH channel were identified manually on the corresponding fluorescent channel. Distance between the DNA-FISH signal and the nucleolar marker immunofluorescence signal was calculated using ImageJ (version 2.14.0/1.54F) and used to count the number of foci contacting nucleolus and the number of cells with at least one contact with the nucleolus.

### Published datasets

Published datasets used in the study were from: H3K27ac-, H3K4me1-, H3K27me3-ChIP-seq ^46^ (GSE72164); H3K4me3- and H3K9me3-ChIPseq ^63^ (GSE23943); H3K9me2-ChIPseq ^64^ (GSE77420); A/B compartments ^65^; Nucleolar DamID and RNAseq in ESCs and NPCs ^19^ (GSE150822); Lamin-DamID ^66^ (GSE17051); SPRITE ^29^ (GSE114242); seqFISH+ ^30^.

## Acknowledgements

This work was supported by the ERC grant (ERC-AdG-787074-NucleolusChromatin), Swiss National Science Foundation (31003B-201268; 320030-227818), and EMBO Postdoctoral Fellowship (ALTF 209-2021) and the Forschungskredit of the University of Zurich to K.W. We thank Catherine Aquino and the Functional Genomic Center Zurich for the assistance in sequenscing. We thank the Scientific Center for Optical and Electron Microscopy of ETH Zurich for support and assistance in LCM. We thank Ana Pombo for her technical advises on laser-microdissection and library preparation and Bing Ren for the F123-ESC line.

## Author contributions

K.W. and J.K-R. established the NoLMseq. K.W. performed NoLMseq in all the described conditions, ChIPseq and RNAseq in F123-ESCs and all data analyses of NADs. S.G. and M.R. performed DNA-FISH analyses. C.M. performed IF analyses. K.W. and R.S. wrote the manuscript. R.S conceived and supervised the project.

## Declaration of interests

The authors declare no competing interests.

## Extended Data figure legends

**Extended Data Figure 1.**
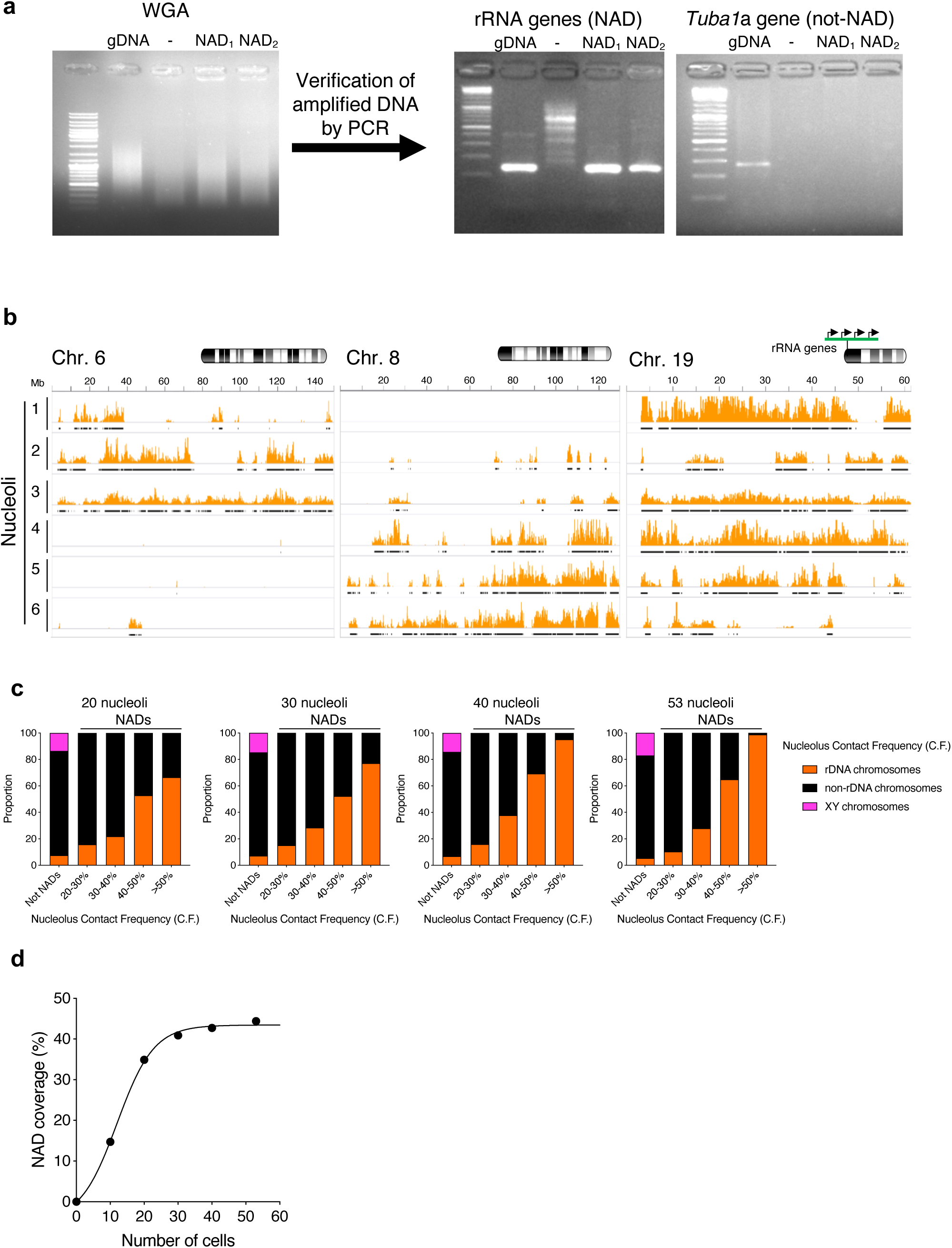
NoLMseq identifies NADs in single cells. **a.** Gel electrophoresis showing whole-genome amplification (WGA) of isolated nucleoli (left panel) and the PCR amplification for rRNA genes and *Tuba1* sequences. **b.** NoLMseq tracks from 6 microdissected nucleoli at chromosomes 6, 8, at 19. Black bars below the tracks indicate NADs called in respective single nucleoli. **c.** Cumulative histogram of genome-wide nucleolar contact frequency (CF) values in 20, 30, 40 randomly selected microdissected nucleoli and all the 53 analysed nucleoli. The location at chromosomes containing rRNA genes (rDNA-chromosomes), without (non-rDNA chromosomes), and X and Y chromosomes is shown. **d.** NAD coverage calculated from 20, 30, 40 randomly selected microdissected nucleoli and all the 53 analysed nucleoli.

**Extended Data Figure 2.**
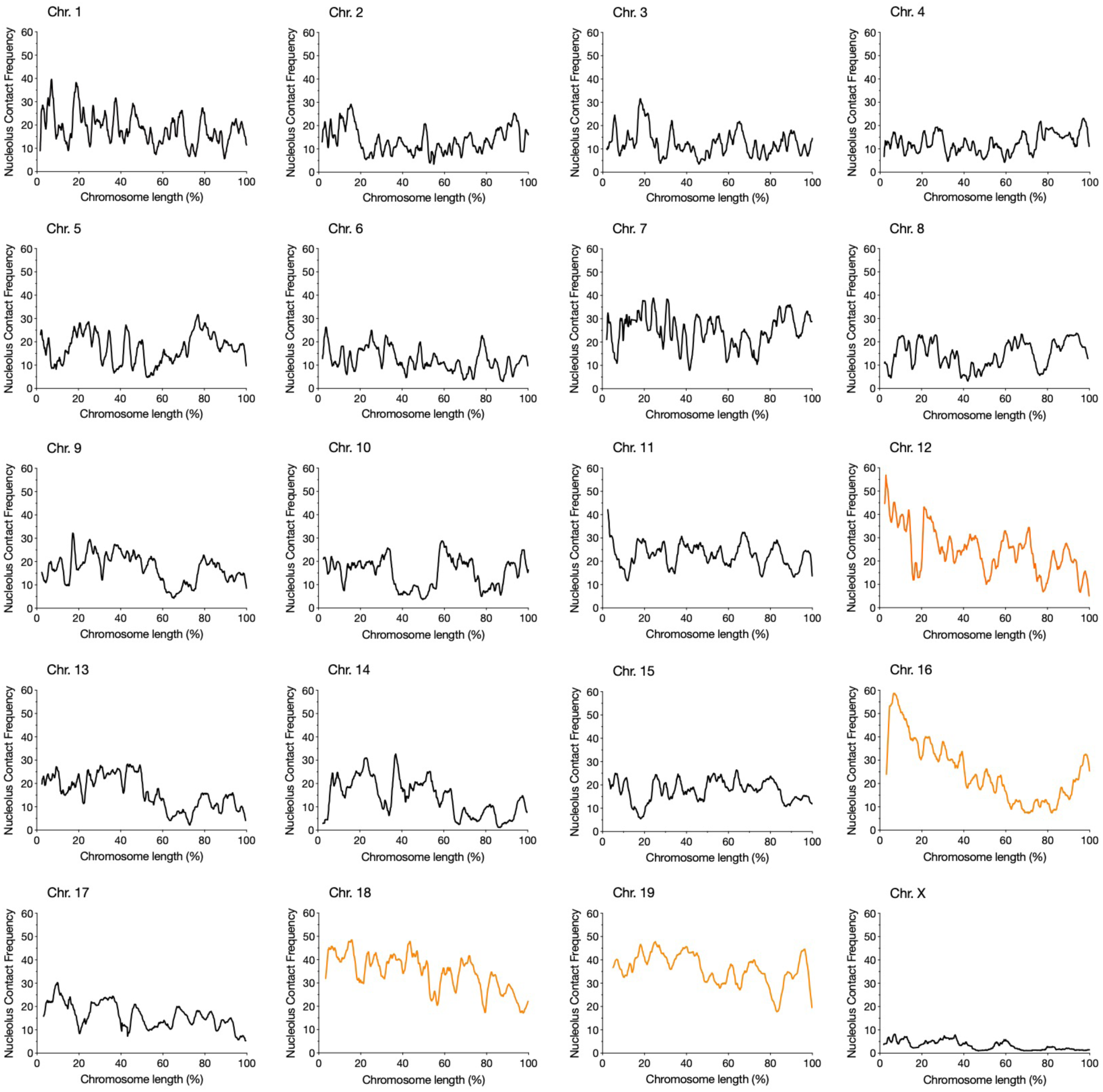
NAD contact frequency for each chromosome. Nucleolus contact frequency for the 53 ESC nucleoli along the length of each chromosome. rDNA-chromosomes are depicted with an orange line.

**Extended Data Figure 3.**
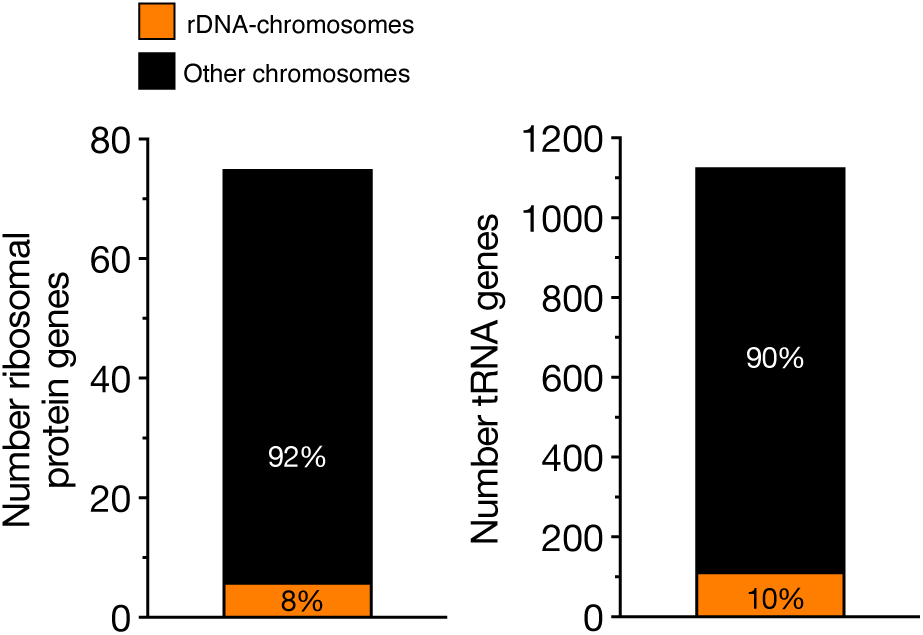
Ribosomal protein and tRNA genes show high nucleolar contact frequency. Distribution of ribosomal protein and tRNA genes at rDNA-chromosomes and chromosomes not containing rRNA genes (other chromosomes).

**Extended Data Figure 4.**
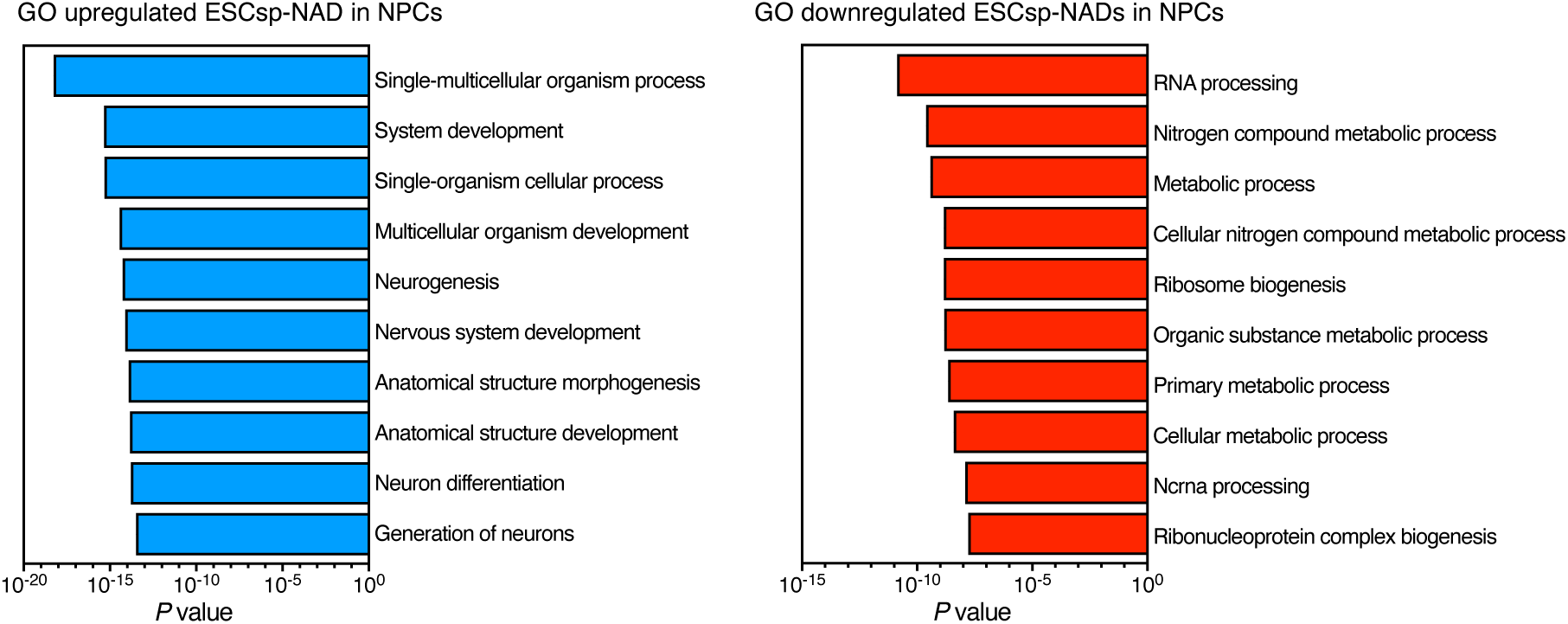
NADs from single nucleoli of neural progenitors. Gene ontology (GO) terms for upregulated and downregulated genes at ESC_sp_-NADs in NPCs.

**Extended Data Figure 5.**
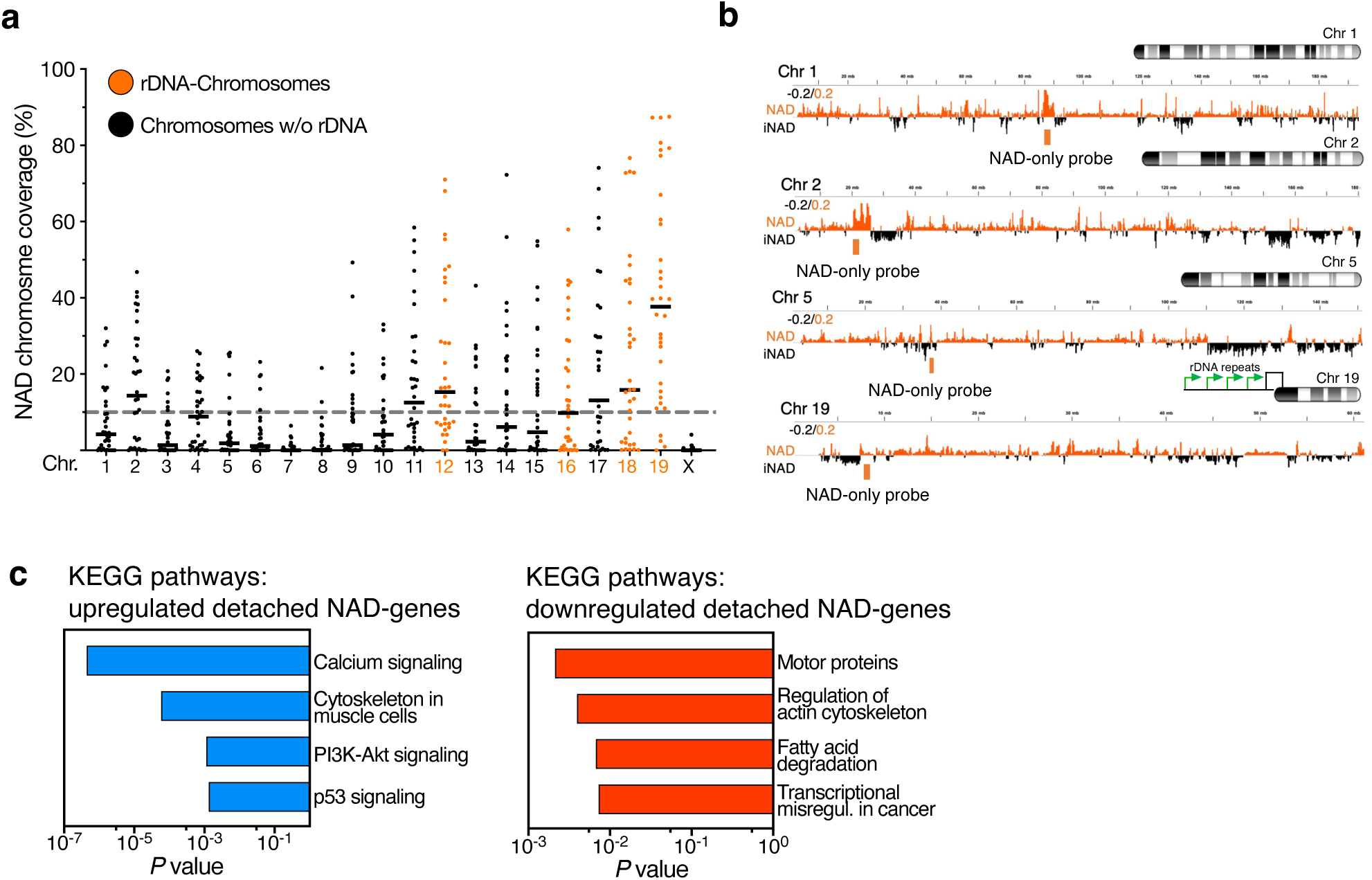
Nucleolar integrity is required for genomic contacts with nucleoli. **a.** Coverage of NADs at each chromosome of single nucleoli from ESCs treated without and with Actinomycin D (ActD). Dotted grey line indicates the average NAD coverage in single cells. Black lines indicate the mean on each chromosome. **b.** Representation of the DNA-FISH probes targeting NADs at chromosomes 1, 2, 5, and 19, the latter containing rRNA genes at the 5’ end, close to the centromeric region. The corresponding NADs profiles from Nucleolar-DamID ^19^ are shown. iNAD: genomic domains not interacting with nucleoli. **c.** KEGG pathways for upregulated and downregulated NAD-genes detached from nucleoli in ESC+ActD.

**Extended Data Figure 6.**
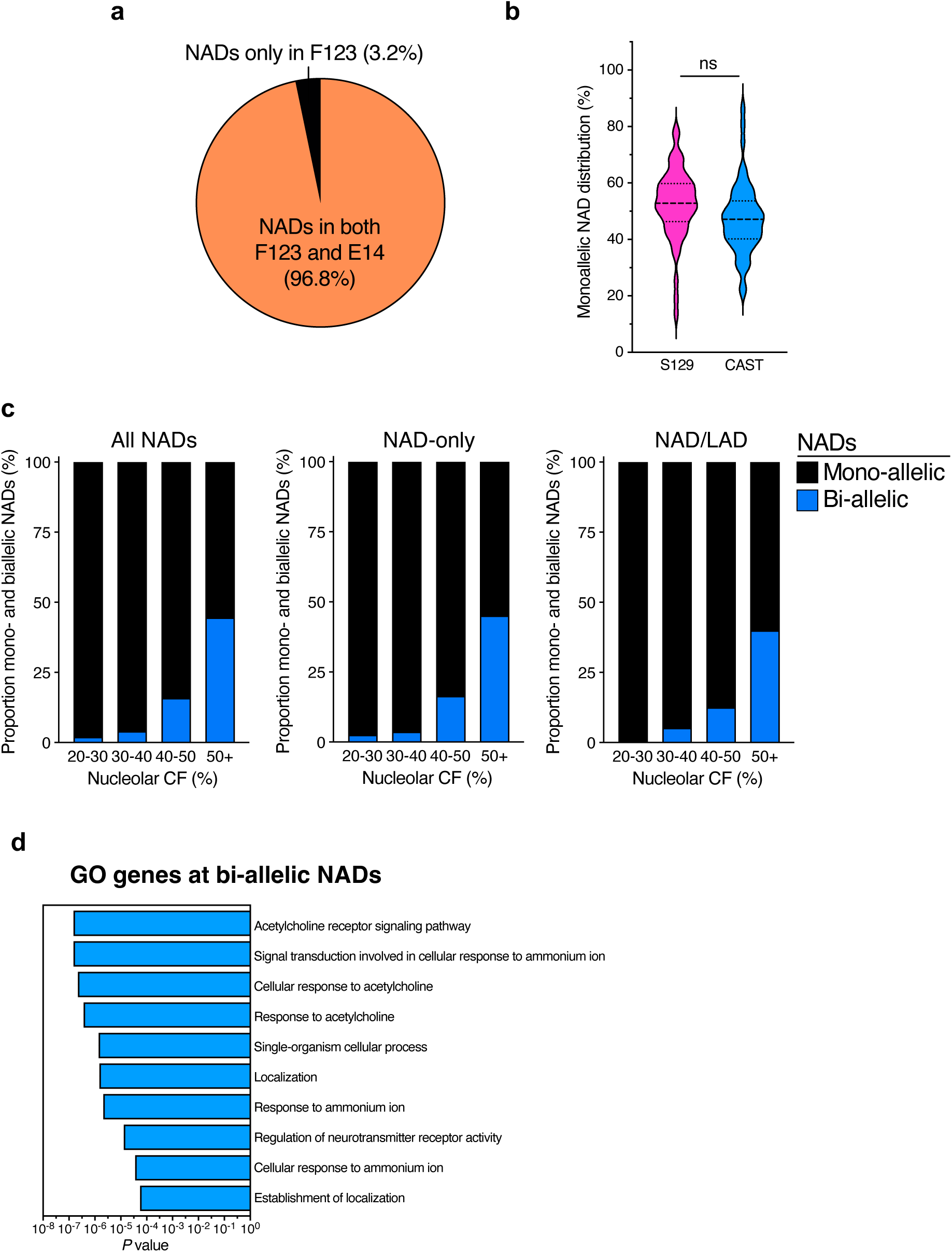
Genomic contacts with nucleoli are mono-allelic, but not parent specific. **a.** Pie chart showing the proportion of NADs identified in F123-ESCs that have also been found in E14-ESCs. **b.** Mono-allelic distribution of 129 and CAST genome around nucleoli of single cells. Black dashed lines indicate median and black dotted lines indicate quartiles. Statistical significance (P-value) was calculated using the unpaired two-tailed t test (ns: non-significant). **c.** Cumulative histogram of nucleolar contact frequency (CF) values with respect to their mono-and biallelic content for all NADs, NAD-only, and NAD/LAD. **d.** Gene ontology (GO) terms of genes located on bi-allelic NADs.

## Notes

### Competing Interest Statement

The authors have declared no competing interest.

### Summary of Updates

This version of the manuscript has been revised to update text and Suppl. Figs. 5,6

